# Evidence for reduced choroid plexus volume in the aged brain

**DOI:** 10.1101/2024.12.11.627733

**Authors:** R Youh, C Perera, IF Harrison, MF Lythgoe, DK Wright, S Nizari, JA Wells

## Abstract

**Background:** The choroid plexus plays an important role in brain homeostasis, including the active secretion of cerebrospinal fluid. Its function and structure have been reported to be affected by normal ageing. However, existing measures of choroid plexus volume may be complicated by partial volume (*in vivo* MRI) and tissue fixation artefacts (histology). In this study, we investigate possible changes in choroid plexus volume within the lateral ventricles of aged mice utilising two structural MRI protocols explicitly designed for time-efficient, high-resolution *in vivo* imaging of the choroid plexus.

**Methods:** Two MRI sequences were utilised to examine *in vivo* choroid plexus volume in the lateral ventricles of young (∼6 months) and aged (∼24 months) mouse brains: 1) an ultra-long echo-time T2 weighted fast-spin-echo and 2) a multi-TE T2* mapping protocol. A test-retest study was performed on a subset of the data to examine the reproducibility of choroid plexus volume estimation. A two-way ANOVA test was performed to determine possible differences in choroid plexus volume in young and aged mouse groups across the two distinct MRI protocols.

**Results:** Reproducibility tests showed a low test-retest variability of the manual segmentation pipeline for both MRI protocols. A statistically significant reduction of *in vivo* choroid plexus volume was found in the aged mouse brain. This finding is concordant with previous histology studies that have observed a reduction in epithelial cell height with ageing across a wide range of species.

**Conclusions:** We present an *in vivo* investigation of changes to lateral ventricle choroid plexus volume in the mouse brain utilising a manual segmentation approach based on two bespoke MRI protocols designed for time-efficient high resolution imaging of the choroid plexus. Furthermore, based on these protocols, we provide evidence for a reduction in choroid plexus volume in the aged brain. This research provides insight for studies utilising MRI measurements of choroid plexus volume as a biomarker of age-related neurologic conditions as it indicates that the ageing process itself does not result in hypertrophy of the choroid plexus, but a decrease in tissue volume.

## Introduction

The choroid plexus (ChP) resides in the lateral, third, and fourth ventricles of the brain and forms the blood-cerebrospinal fluid barrier (BCSFB). The ChP performs several key roles in service of normal brain function including the active secretion of cerebrospinal fluid (CSF) which may in turn support the efficacy of CSF-mediated brain-clearance mechanisms such as the glymphatic pathway [1, 2]. Given this multifaceted and unique physiology, the ChP has been proposed as a hitherto underexplored site of mechanistic significance in age-related conditions such as multiple sclerosis and Alzheimer’s disease [3-6].

Towards the goal of developing novel imaging biomarkers of ChP derangement, there has been an emergence of publications reporting changes in ChP volume associated with disease, based on segmentation of high-resolution structural MRI scans [7-15] . One notable observation in the context of age-related neurodegenerative disease, are reports increased lateral ventricle (LV) ChP volume in the aged human brain [16, 17]. Interestingly, this finding would appear to contradict direct histological assessment where ChP epithelial atrophy has been reported with age in both mice and rats as well as in the human brain [18-20]. These reports are further complicated by possible methodological limitations associated with both approaches such as partial volume effects exacerbated by the retrospective analysis of structural MRI scans that were not explicitly designed for reliable ChP segmentation as well as possible artefacts due to tissue fixation procedures required in histology. Thus, we aimed to help disambiguate the relationship between LV ChP volume and ageing by acquiring *in vivo* measurements of ChP volume using two bespoke time-efficient MRI protocols designed for high-resolution *in vivo* imaging of the ChP: i) an ultra-long echo-time (TE) T2 weighted fast-spin echo (FSE) scan and ii) a multi-TE T2* mapping protocol. We demonstrate that both MRI protocols provide measures of choroid plexus volume with low test-retest manual-segmentation variability. We then apply both imaging protocols to aged and young-adult control mice for non-invasive, *in vivo* estimation of ChP volume. Doing so returns evidence for a reduction in choroid plexus volume in the aged brain, suggesting that the ageing process *per se* does not lead to an increased volume of ChP tissue, but a hypotrophic decrease.

## Methods

### Animal preparation

All experiments were conducted in accordance with the European Commission Directive 86/609/EEC (European Convention for the Protection of Vertebrate Animals Used for Experimental and Other Scientific Purposes) and the United Kingdom’s Home Office Animals (Scientific Procedures) Act (1986). Prior to data collection, mice were acclimatized in an animal facility with a 12-hour light/12-hour dark cycle, and food and water were provided ad libitum.

Imaging was performed using a horizontal-bore 9.4T Bruker preclinical system (BioSpec 94/20 USR; Bruker) equipped with a 440-mT/m gradient set featuring outer and inner diameters of 205 mm and 116 mm, respectively (BioSpec B-GA 12S2). An 86-mm volume transit RF coil and a four-channel receiver-array coil, specifically designed for mouse brain imaging (Bruker), were used for data acquisition. Mice were anesthetized with a 2% isoflurane mixture (4:1 room air/O_2_), adjusted to 1.5% to maintain a respiratory rate of approximately 150 bpm, which was continuously monitored using a pressure pad throughout the scan. The mouse’s head was rigidly fixed using blunt ear bars. Core body temperature was measured with a rectal probe (SA Instruments, Stony Brook, NY) and kept at 37.0 ± 0.5°C, regulated by an adjustable water bath connected to a mouse heating pad (Bruker BioSpec; Bruker, Kontich, Belgium). MRI experiments were performed on male C57BL/6JRj mice at 6 and 24 months of age (Janvier labs) (n = 10 and n = 12 respectively). This corresponds to an age in human years of approximately 30 and 70 years respectively.

### Data acquisition

In this study, two structural MRI protocols designed for high resolution/contrast imaging of mouse ChP were applied in separate scan sessions: (1) 3D FSE ultra-long TE readout with the following parameters: FOV = 19.6 x 19.6 x 3.6mm (centred around the LVs,-see Figure 1); matrix size = 196 x 196 x 36; 0.1mm isotropic resolution; echo train length = 64; effective TE = 176.2ms; TR = 5000ms; 1 average; acquisition time ∼6.5 minutes; (2) 3D multi-TE gradient echo with the following parameters; FOV = 16 x 19.25 x 12 mm; matrix size = 128 x 154 x 96; TEs = 2.19, 5,25, 8.31, 11.37, 14.43, 17.49, 20.55, 23.61, 26.67, 29.73, 32.79, 35.85 ms; TR =66ms; flip angle = 15°;1 average; 0.125mm isotropic resolution; acquisition time ∼ 16minutes.

**Figure 1.**
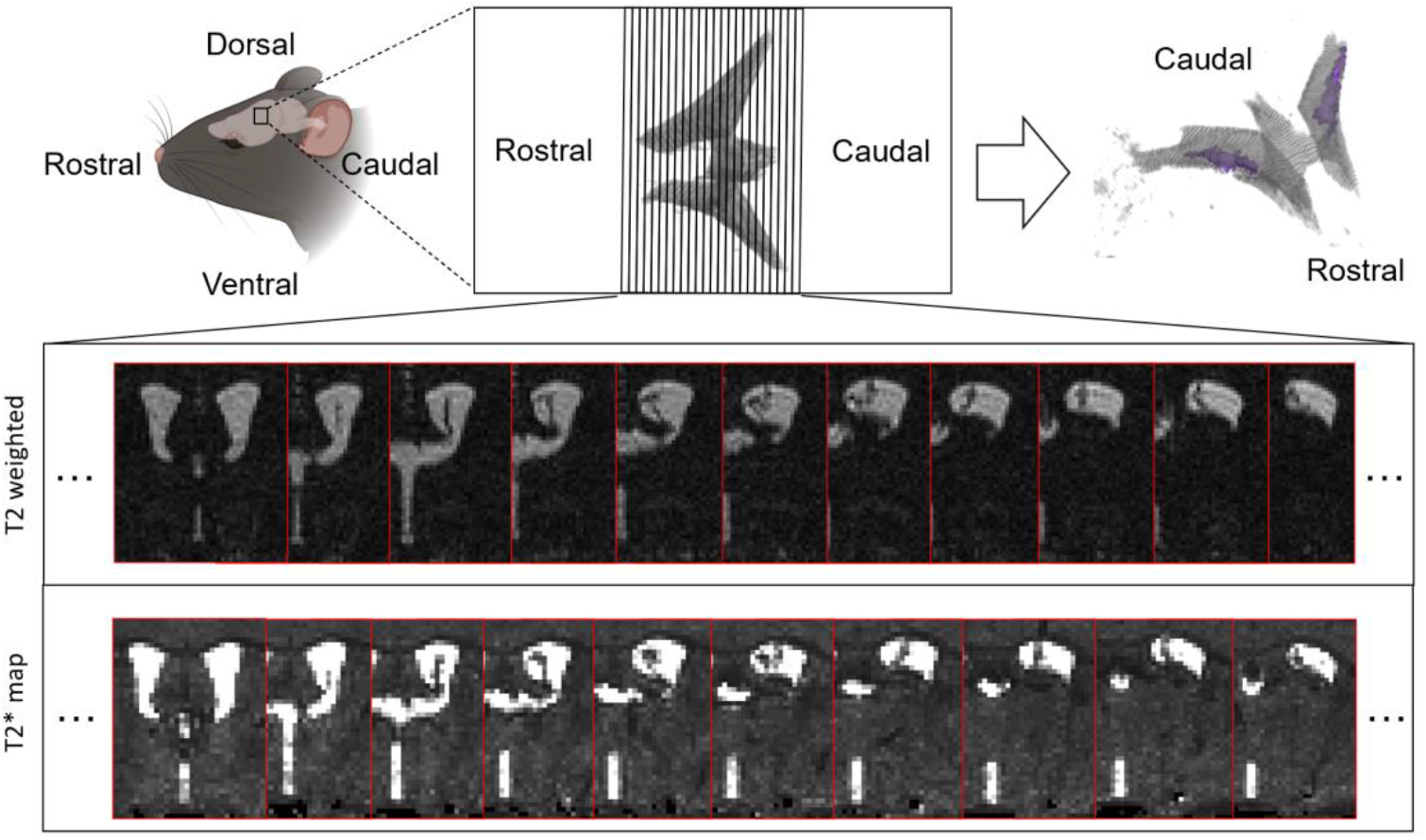
Schematic of manual segmentation pipeline. Top: The ChP in this figure (upper right) is labelled within the LV and is highlighted in purple. Bottom: Representative datasets from the same subject utilising two different imaging protocols (T2w and T2* map, n = 1 representative), which demonstrates a collection of sequential slices that capture the LV for ChP volume estimation.

Both imaging datasets were collected to estimate the volume of ChP. The raw MRI images of the 3D FSE h-T2w readout were directly utilised, while the 3D multi-TE GRE T2*w readout were then used to generate T2* map images for ChP volume estimation by fitting the data to a simple mono-exponential model using Matlab (Mathsworks, Massachusetts, USA).

### ChP volume estimation

A manual segmentation pipeline was conducted for *in vivo* ChP volume estimation in young and aged mouse brains among all image datasets (Figure 1). For each slice, the ChP was manually segmented based on visual inspection of the T2w images/ T2* maps. The segmentations were carried out using the Volume Segmenter toolbox in MATLAB. The manual segmentation was performed blind to the animal group (aged vs. young control). To investigate the reproducibility of ChP volume estimation using the manual segmentation pipeline, the segmentation pipeline was repeated on 3 randomly selected subjects within a one-day interval. This was performed separately for both MRI protocols. The mean absolute % difference in volume measurements was calculated for each subject.

### Statistical analysis

The effect of age and imaging protocol (ultra-long TE T2w & T2*maps) on ChP volume was investigated using a two-way analysis of variance (ANOVA) test using GraphPad Prism (GraphPad Software, Boston, Massachusetts USA). Errors are reported as standard deviations from the mean.

## Results

### ChP Segmentation Reproducibility test

Reproducibility tests were carried out to investigate the consistency of the manual ChP volume segmentation methods (Figure 2). An average absolute change of 9.38% was observed between the repeatedly segmented datasets using high-resolution ultra-long TE T2w MRI images (Figure 2 (A)), while an average absolute change of 7.07% was observed using T2* map images (Figure 2 (B)). A representative example of the reproducibility test is shown in Figure 3(C)(D), where we observe a high degree of segmentation agreement for both protocols. Here, we established that the manual segmentation method yielded reproducible estimates of ChP volume.

**Figure 2.**
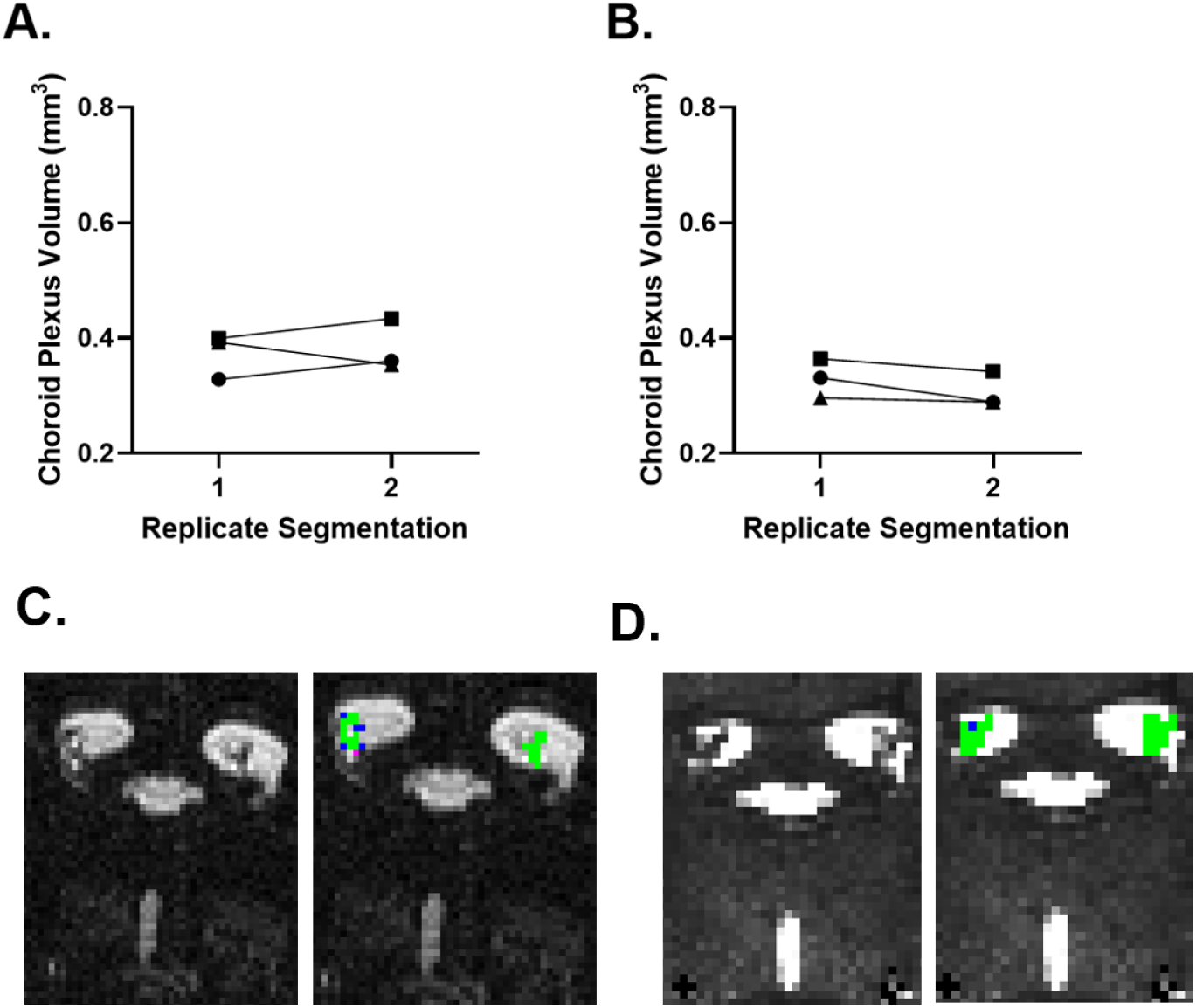
Segmentation pipeline reproducibility tests. (A) ChP volume segmentation reproducibility test based on T2w datasets. (n=3, randomly selected) (B) ChP volume segmentation reproducibility test based on T2* map datasets. (n=3, randomly selected) (C) Representative example of visual comparison based on a T2w MRI image obtained from the reproducibility test. (D) Representative example of visual comparison based on a T2* map image obtained from the reproducibility test. For (C) and (D), the green areas indicate the regions segmented on both the first and second days. The purple areas represent the segmentations made on the first day that were not included on the second day. Conversely, the blue areas represent the segmentations made on the second day that were absent on the first day.

**Figure 3.**
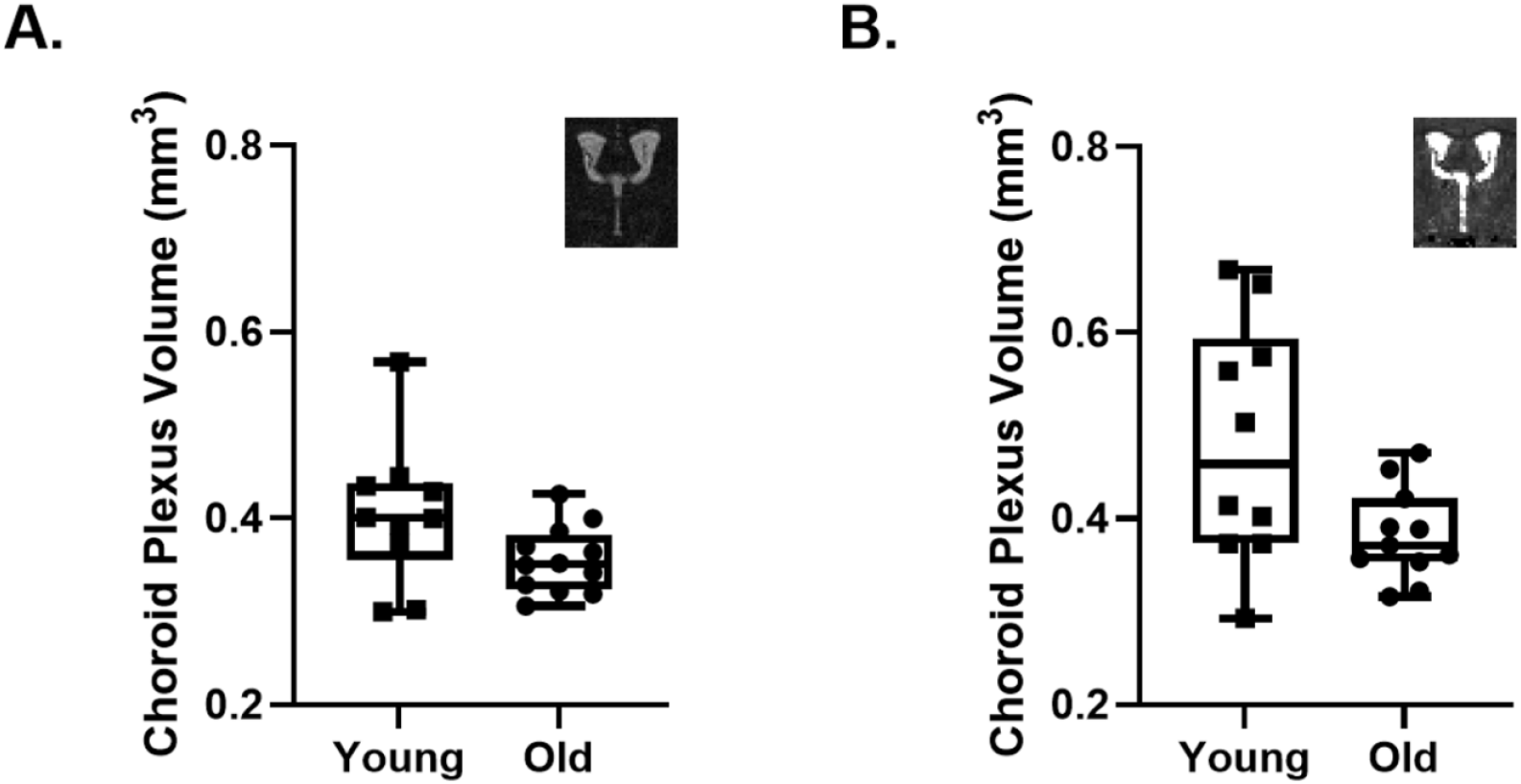
Comparison of the ChP volume estimated in young and old groups from high-resolution T2-weighted MRI images (A) and T2* map MRI images (B). Each dot represents an individual mouse.

### *In vivo* choroid plexus volume in the aged mouse brain

A two-way ANOVA test was conducted to compare the main effects of age and MRI protocol as well as the interaction effects on the ChP volume. Prior to unblinding, one mouse was excluded from all T2* map analyses due to MRI artifacts within the LV that affected the segmentation. No other animals were excluded from the analysis. The age effect was statistically significant (*P* < 0.01) with a relatively large effect size (*ω*^2^ > 0.14). The effect of imaging datasets was statistically significant (*P* < 0.05) with a medium effect size (0.06 < *ω*^2^ < 0.14). The interaction effects showed no significance (*P* > 0.05). A comparison of ChP volume between young (*N* = 10, 0.405 ± 0.076*mm*^3^) and aged mice (*N* = 12, 0.356 ± 0.036 *mm*^3^) from ultra-long TE T2-weighted MRI images is shown in Figure 3(A). A comparison of ChP volume between young (*N* = 10, 0.481 ± 0.129 *mm*^3^) and aged mice (*N* = 11, 0.383 ± 0.050 *mm*^3^) from T2* map images is shown in Figure 3(B). These findings indicate a statistically significant difference in ChP volume between the young and old groups.

## Discussion

In this study we applied two time-efficient MRI protocols designed for robust estimation of ChP volume in the LVs of the mouse brain. The protocols employed were an ultra-long TE FSE and a multi-TE gradient echo T2* mapping sequence. We then evaluated the reproducibility of choroid plexus volume estimates from manual segmentation of the images which returned a mean absolute difference of 9% and 7% for each protocol respectively, providing a solid foundation to detect putative group-wise differences in ChP volume in mouse models. We then applied the protocols to a cohort of aged and young mice to investigate possible changes in ChP volume in the ageing brain, given conflicting reports in the literature. Doing so, we found evidence for reduced ChP volume in the aged mouse brain, a finding concordant with observations of ChP epithelial atrophy from direct histological assessment [18-20]. This finding suggests that ageing causes atrophy of the choroid plexus and that this can be readily detected using non-invasive *in vivo* structural MRI.

The concept of the ultra-long FSE sequence for time-efficient imaging of ChP volume is a simple one that has been used before in MRI studies primarily designed to image the CSF [21, 22]. This employs a long echo train (in this case 64) which allows the acquisition of multiple lines of k-space after a single excitation for time-efficient data capture (in this case giving 0.1mm isotropic resolution images in ∼6 minutes). The resultant long echo time (TEeff = 176ms) will highly attenuate the signal from all tissue types within the brain with the exception of CSF due to its uniquely long T2 relaxation time. Thus, in these images, negative contrast is employed where the CSF appears bright and the ChP tissue is dark. Accordingly, a downside of this approach is that all extra-ventricular tissue signal is nulled in these images meaning that this protocol will only inform about ventricular and ChP structure and not other grey/white matter regions for example. Here we implemented a 3D FSE sequence as image quality was found to be poor using an equivalent 2D sequence with matched geometry likely due to the cumulative effect of multiple refocusing pulses when using 0.1mm slice thickness (data not shown). The multi-echo gradient echo 3D sequence is designed for time-efficient T2* mapping using a short TR and reduced flip angle. Similarly, this exploits the marked difference in T2* between the ChP and adjacent CSF. Unlike the ultra-long TE FSE sequence however, this also yields T2* maps across the extra-ventricular tissue which could be useful for detection of other radiological markers of pathology such as micro-hemorrhages for example.

In regards to employing a more automated and objective approach to ChP volume estimation, within the ultra-long TE FSE data we explored using signal-intensity based thresholding across the lateral ventricles under the premise that there would be two distinct signal populations at ∼0 signal (ChP) and >> 0 signal (CSF). However, the distribution of signal intensities within the LVs was found to more continuous and not as bi-modal as we initially surmised meaning that it was challenging to define a ‘cut-off’ signal intensity to accurately separate ChP and CSF voxels. We suspect that this principally reflects the partial volume of the CSF with tissue at the edge of the ventricles and ChP in addition to some subtle Gibbs ringing artefacts in the CSF (whose characteristic pattern can be distinguished by eye). Thus, we proceeded with a manual segmentation approach which further benefits from the experience of the rater to recognise the characteristic morphology of the ChP within the LVs (see Figure 1). Indeed, this approach returned a mean absolute error in ChP volume estimation of 9% (ultra-long TE FSE) and 7% (multi-TE GE T2* maps) following test-retest ChP segmentations suggesting that this approach can yield reliable estimates of ChP volume. Moving forward, the use of more automated approaches may alleviate the need for the time-consuming manual segmentation [17, 23, 24]. Interestingly, the estimates of ChP volume based on the T2* maps were greater than those using the ultra-long TE FSE protocol which likely stems from a ‘blurring’ of the ChP tissue due to the macro time-invariant susceptibility effects that are present with T2* vs T2 contrast.

Our finding of reduced ChP volume in the aged brain is consistent with histological observations of epithelial cell atrophy in the aged mouse, rat and human brain [18-20]. The precise reasons why, using *in vivo* MRI structural MRI scans, we find a reduction in ChP volume in the aged mouse brain whereas others have reported an increase in ChP volume with ageing in the human brain remains unknown at this time. Here, we speculate the following methodological/physiological factors may be at play: i) the limited spatial resolution of *in vivo* structural MRI scans to accurately spatially resolve the fine structure of the ChP may lead to systematic errors that influence the accuracy of ChP volume measurement. This may be impacted by changes in the volume of the LVs that occur with ageing, in turn altering the expanse of fluid in which the ChP tissue can float within. ChP tissue that is diffusely present in a larger volume of ventricular CSF could be misinterpreted as possessing a greater volume then ChP tissue more densely packed into a small volume of lateral ventricular CSF due to partial volume effects given the limited spatial resolution of the images. In the mice examined here, no significant differences in our *in vivo* measurements of ventricular volume were detected between the aged and young adult groups, as found in our previous study [25] (data not shown). ii) earlier studies have reported an increased volume of ChP tissue in a range of neurological conditions (e.g. [11, 26-30]). Of note, to our knowledge a decrease in ChP volume associated with a disease has yet to be reported. Therefore, the increased ChP volume observed in the aged human brain may reflect undiagnosed comorbidities with human ageing that are not necessarily recapitulated in the aged mouse brain, highlighting the limitations of the mouse to model the complex multifaceted co-morbidities that come with human ageing. iii) the ChP is known to undergo marked movement due to cardiac and respiratory pulsation. As, to our knowledge, none of the previous studies of ChP volume have been gated to cardiac or respiratory signals (including the present work), this movement may introduce a degree of inaccuracy into the structural images where ChP tissue that undergoes a greater degree of motion presents with a spurious increase in volume due to blurring during the acquisition (which typically takes several minutes). This possible artefact may have an age dependence where, in the case of cardiac pulsation, upstream stiffening of the vessels leads to greater pulse wave propagation to the downstream cerebral vessels [31]. Our preliminary investigation into the degree of ChP motion with cardiac and respiratory cycles suggested that this motion was subtle however and will thus have relatively little influence on our ChP volume measurements (data not shown).

As described above, previous research has reported the association between ChP volume and neurological disorders such as Alzheimer’s disease (AD) and multiple sclerosis (MS). An investigation found that an increased ChP volume is associated with patients with AD, as determined by T1w MRI scans [30]. This study also emphasizes the need for more accurate measurement techniques to delineate ChP volume in MRI scans, alongside the development of effective segmentation methods [ref1]. In addition, a retrospective analysis indicated that the ChP volume within the LV is greater in individuals with AD compared to those experiencing subjective cognitive impairment (SCI) or mild cognitive impairment (MCI), as assessed through 3D T1-weighted (T1w) imaging [15]. Apart from AD, a previous study has shown that patients with MS, particularly those with the relapsing-remitting form, exhibit a larger ChP volume compared to healthy individuals based on 3D T1w images and that this enlargement may be partially related to inflammatory processes [32].

To conclude, in this work we aimed to better understand the impact of ageing on ChP volume, captured using simple, translational, structural MRI scans. We did this by applying two MRI protocols specifically designed for efficient, high resolution, high contrast imaging of the ChP, to the young and aged mouse brain. We find evidence for a reduction in ChP volume in the aged brain, with both methods returning similar findings. This provides evidence that ageing itself results in atrophy and not hypertrophy of the ChP.

## Acknowledgements

JAW, CP and SN are supported by the Wellcome Trust (225345/Z/22/Z). DW is supported by the National Health and Medical Research Council to DKW [grant number: 1174040]. IFH is supported by both Parkinson’s UK (F-1902) and Alzheimer’s Research UK (ARUK-RF2019A-003). MFL is supported by Rosetrees Trust and the John Black Charitable Foundation (Grant No. A2200); Medical Research Council (MR/M009092/1); the Brain Tumour Charity (Grant No. QfC_2018_10387); the Edinburgh-UCL CRUK Brain Tumour Centre of Excellence (Grant No. C7893/A27590); the CRUK & EPSRC Comprehensive Cancer Imaging Centre at KCL and UCL (Grant No. C1519/A16463 and C1519/A10331).

